# STing: accurate and ultrafast genomic profiling with exact sequence matches

**DOI:** 10.1101/855478

**Authors:** Hector F. Espitia-Navarro, Aroon T. Chande, Shashwat D. Nagar, Heather Smith, I. King Jordan, Lavanya Rishishwar

**Affiliations:** School of Biological Sciences, Georgia Institute of Technology, Atlanta, GA 30332, USA; PanAmerican Bioinformatics Institute, Cali, Valle del Cauca 760043, Colombia; Applied Bioinformatics Laboratory, Atlanta, GA 30332, USA; School of Mathematics, Georgia Institute of Technology, Atlanta, GA 30332, USA

## Abstract

Genome-enabled approaches to molecular epidemiology have become essential to public health agencies and the microbial research community. We developed the algorithm STing to provide turn-key solutions for molecular typing and gene detection directly from next-generation sequence data of microbial pathogens. Our implementation of STing uses an innovative *k*-mer search strategy that eliminates the computational overhead associated with the time consuming steps of quality control, assembly, and alignment required by more traditional methods. We compared STing to six of the most widely used programs for genome-based molecular typing and demonstrate its ease of use, accuracy, speed, and efficiency. STing shows superior accuracy and performance for standard multilocus sequence typing schemes, along with larger genome-scale typing schemes, and it enables rapid automated detection of antimicrobial resistance and virulence factor genes. We hope that the adoption of STing will help to democratize microbial genomics and thereby maximize its benefit for public health.

## Main

Molecular typing entails the identification of distinct evolutionary lineages (i.e. types) within species of bacterial pathogens; it is an essential element of both outbreak investigation and routine infectious disease surveillance^1, 2^. Multilocus sequence typing (MLST) was developed as the first sequence-based approach to molecular typing in 1998^3^. Initially, MLST schemes relied on Sanger sequencing of PCR amplicons from fragments of 7-9 housekeeping genes spread throughout the genome. While this approach truly revolutionized molecular epidemiology, it is time consuming and costly compared to current next-generation sequencing (NGS) methods. Nevertheless, MLST remains widely used for molecular typing, particularly in light of valuable legacy data relating sequence types (STs) to epidemiological information.

Public health agencies increasingly couple NGS characterization of microbial genomes with downstream bioinformatics analysis methods to perform molecular typing. The overhead associated with the bioinformatics methods used for this purpose, in terms of both the required human expertise and computational resources, represents a critical bottleneck that continues to limit the potential impact of microbial genomics on public health. This is particularly true for local public health agency laboratories, which are typically staffed with microbiologists who may not have substantial bioinformatics expertise or ready access to high-performance computational resources. In light of this ongoing challenge, our group is working to develop turn-key solutions for genome-enabled molecular epidemiology, including both molecular typing and the detection of critical antimicrobial resistance (AMR) and virulence factor (VF) genes. Methods of this kind must be easy to use, computationally efficient, fast, and most importantly, highly accurate.

We previously developed stringMLST as an alternative approach to genome-enabled molecular typing of bacterial pathogens^4^. stringMLST relied on *k*-mer matching between NGS sequence reads and a database of MLST allele sequences, thereby eliminating the need for the sequence quality control, genome assembly, and alignment steps that the first generation of genome-enabled typing algorithms used. It proved to be accurate and fast for traditional MLST schemes, but it did not scale well to the larger genome-scale typing schemes, such as ribosomal MLST (rMLST) or core-genome MLST (cgMLST), which are increasingly used in molecular epidemiology^1, 5^. Here, we present our new approach to this problem – STing. The STing algorithm is distinguished from its predecessor in several important ways: the efficiency of its code base, the underlying data structure that is uses, and the scope of its applications. These innovations provide for superior accuracy and performance compared to both stringMLST and other widely used programs for genome-enabled molecular typing. Below, we provide a high-level overview of the STing algorithm, details of which can be found in the Online Methods, and we report on its use across several typing schemes and for automated gene detection.

The STing algorithm breaks down (*k*-merizes) NGS reads into *k*-mers and then compares read *k*-mers against an indexed reference sequence database (Figure 1). The speed and efficiency of the algorithm are derived from the nature of the *k*-mer search strategy used along with the structure of the reference sequence database. For each individual read, a single central *k*-mer is initially compared against the sequence database. Reads are only fully *k*-merized if there is an initial match between the central *k*-mer and the database. If there is no match, which occurs for the vast majority of reads, the read is discarded. This results in substantial savings in terms of both the number of reads that need to be *k*-merized and the number of database search steps. The reference sequence database is indexed as an enhanced suffix array (ESA)^6^; this enables the efficient representation of entire sequences, as opposed to other *k*-mer based methods that employ *k*-merized sequences in hash tables. The ESA data structure allows for a single sequence index, independent of *k*-mer size, whereas the hash table approach necessitates independent indices for each *k*-mer size. Finally, the ESA data structure facilitates rapid exact *k*-mer matches between input reads and the indexed database.

**Figure 1.**
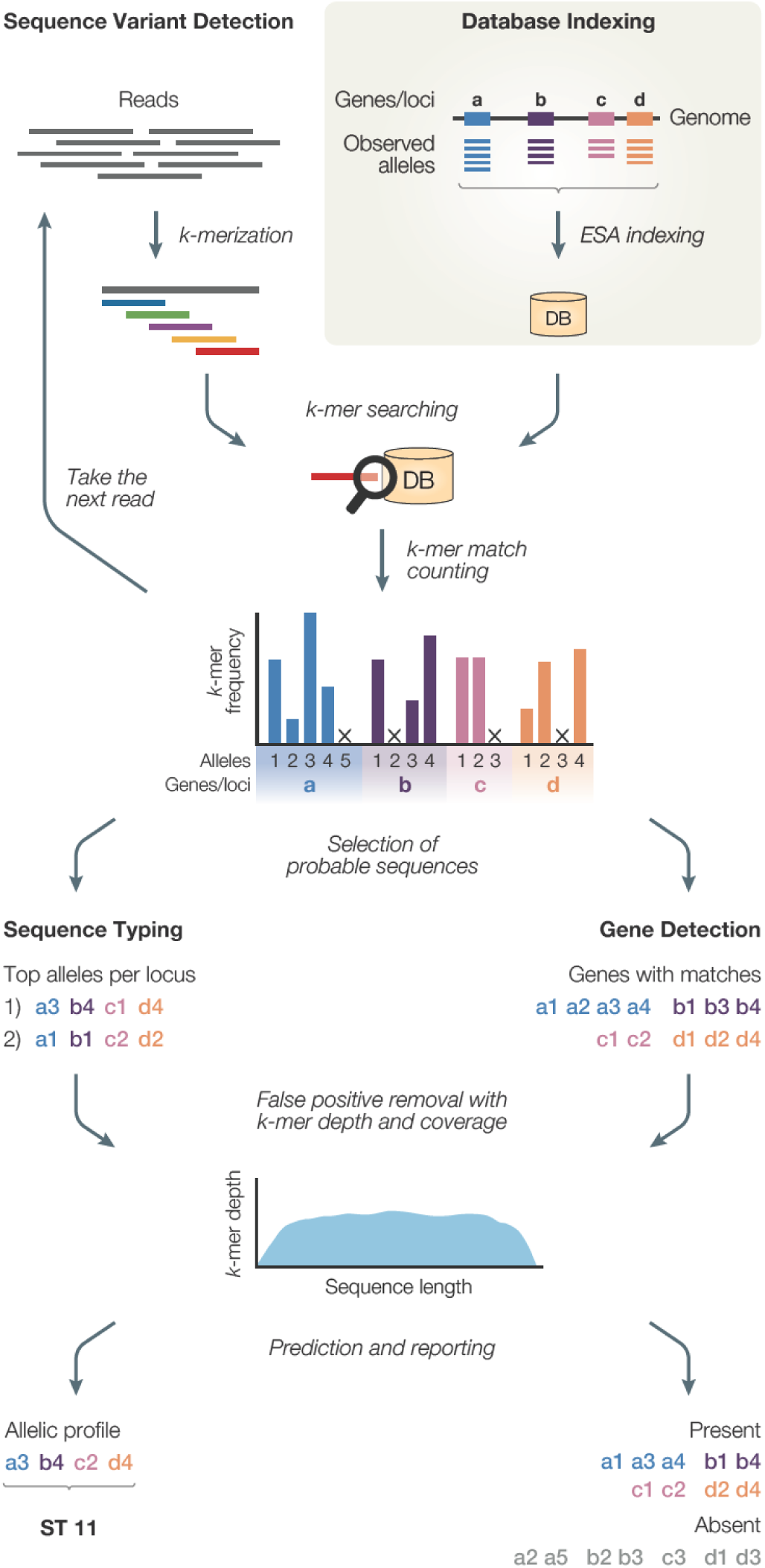
Schematic representation of the STing algorithm. The STing algorithm comprises two main phases: Database indexing (shaded box) – user supplied reference sequences (allele or gene sequences) are transformed into an enhanced suffix array (ESA) index for rapid *k*-mer search during the sequence variant detection phase; and Sequence variant detection – reads are *k*-merized and each *k*-mer is searched within the database. For each match located in the database, a table of frequencies is maintained for the matched sequence within the database. These frequencies are then utilized to select candidate alleles/genes to be present in the samples analyzed. False positive alleles/genes are filtered out by calculating and analyzing *k*-mer depth and sequence length coverage from the selected candidate sequences. Lastly, predictions of allelic profile and ST, and presence/absence of genes, are made and reported. A more detailed flowchart of the algorithm can be seen in Supplementary Figure 1.

STing can be run in two modes – sequence typing or gene detection – and typing can be run in fast or sensitive modes.

We compared STing to six of the most widely used programs for genome-enabled molecular typing, including its predecessor stringMLST (Figure 2). The programs were evaluated for accuracy in terms of the percentage of correct allele predictions, speed in terms of average run time, and efficiency in terms of average maximum RAM consumption. Genome-enabled typing programs can be classified according to the algorithmic paradigm that they use: *k*-mer only, *k*-mer plus alignment, read-to-genome mapping, mapping with local assembly, and full assembly (see Supplement for more information). STing uses the minimalist *k*-mer only approach. STing was run in the fast and sensitive modes for the traditional housekeeping MLST scheme and two larger-scale typing schemes, rMLST and cgMLST. Allele databases for all three typing schemes were taken from the PubMLST database (https://pubmlst.org/). The STing fast mode uses a *k*-mer matching only strategy, and the sensitive mode includes an additional step whereby false positive matches are excluded based on gaps in the coverage profiles of *k*-mer matches to allele sequences.

**Figure 2.**
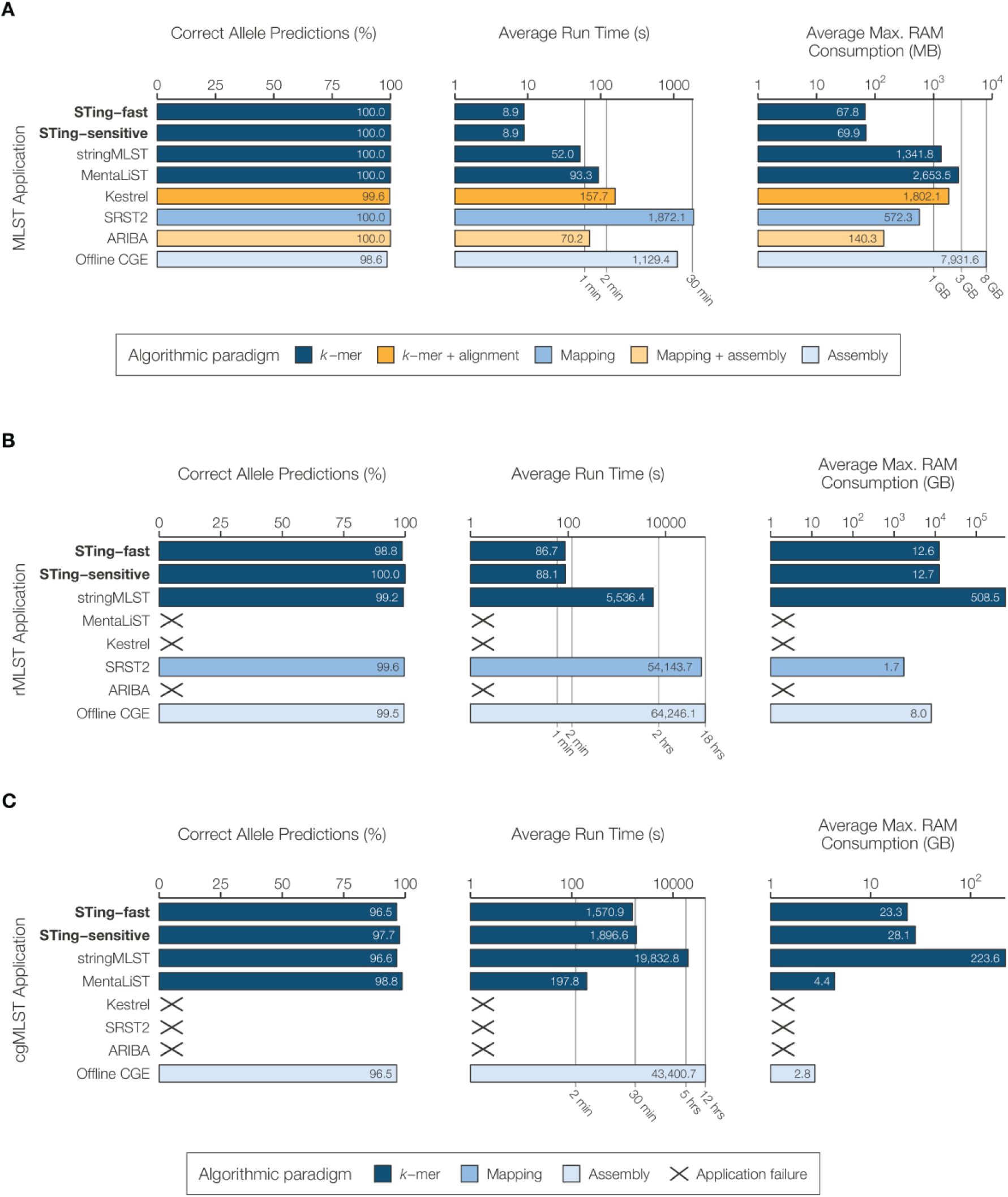
Performance comparison of STing with 6 other sequence typing applications. The fast and sensitive modes of STing are compared to 6 other contemporary typing applications to measure the accuracy and runtime performance, using three different typing schemes: (A) the traditional MLST (loci=7) on 40 samples from four bacterial species (10 samples per species: *C. jejuni, C. trachomatis, N. meningitidis*, and *S. pneumoniae*); (B) the ribosomal MLST (rMLST) scheme (loci=53) on 20 samples of *N. meningitidis*, and (C) the core genome MLST (cgMLST) scheme (loci=1,605) on 20 samples of *N. meningitidis*. The typing applications are color coded based on the algorithmic paradigms that they utilize for performing sequence typing. Performance is measured in terms of the percentage of correct alleles predicted, the average runtime across each dataset measured in seconds (displayed in log-scale), and average peak RAM utilization across each dataset measured in megabytes (MB) for MLST, and gigabytes (GB) for rMLST and cgMLST (both displayed in log-scale).

Comparisons were performed for 10 samples each across four species that are widely used in MLST and accordingly have diverse MLST databases: *Campylobacter jejuni, Chlamydia trachomatis, Neisseria meningitidis*, and *Streptococcus pneumoniae*. STing shows 100% accuracy, in both the fast and sensitive modes, as well as the fastest run time and lowest memory use of any program for MLST (Figure 2A). The results of the same comparisons are broken down for each of the four individual species in Supplementary Figure 2. We also ran STing for MLST across a range of sequence coverage levels in an effort to assess its detection limits and multi-core performance (Supplementary Figure 3). STing performs best at 40x coverage, but it maintains accuracy at 20x with a marginal drop-off at 10x. While STing is designed as a single core application, we found that executing multiple threads of the program allows it to maintain run time up to 40x coverage. This provides for a straightforward way to run STing on numerous genome samples; the MLST accuracy and speed metrics for STing run on a larger dataset of 1,000 *N. meningitidis* samples are shown in Supplementary Table 1. When this large scale analysis was performed, STing was able to uncover seven samples that were initially scored as erroneous predictions but actually turned out to be mis-annotated on the PubMLST database (Supplementary Table 2).

STing also shows the highest accuracy, speed, and efficiency for the four programs that are capable of genome-enabled rMLST typing (Figure 2B). Programs that show as ‘X’ in these comparisons were unable to run for a variety of reasons related to their initial design, the runtime, and database indexing limitations. The program MentaLiST shows marginally higher accuracy, run time, and efficiency for cgMLST compared to STing, which shows the second best metrics for these categories (Figure 2C). However, the utility of MentaLiST, which was designed specifically for cgMLST, is limited by the size of the database that can be indexed. For that reason, it could not be run on the latest rMLST database available from PubMLST.

In addition to molecular sequence typing, STing can also be used for automated gene detection directly from NGS reads. The gene detection mode uses a database of genes of interest, and we used databases of AMR and VF genes given their public health relevance. The Comprehensive Antibiotic Resistance Database (CARD https://card.mcmaster.ca/) of 1,434 AMR genes and the Virulence Factors of Pathogenic Bacteria database (VFDB http://www.mgc.ac.cn/VFs/) of 1,443 VF genes were used for this purpose^7, 8^. STing was used to query the AMR and VF databases with 71 NGS genome datasets for 25 bacterial pathogen species taken from the World Health Organization (WHO) global priority list of antibiotic-resistant bacteria^9^. STing shows very high accuracy metrics for both AMR and VF detection (Figure 3A), along with fast and efficient performance (Figure 3B). STing can be run in in this way to rapidly detect any genes of interest, which extends its utility beyond public health genomics. This could be particularly useful for large scale environmental genomics samples, including amplicon-based and metagenome studies.

**Figure 3.**
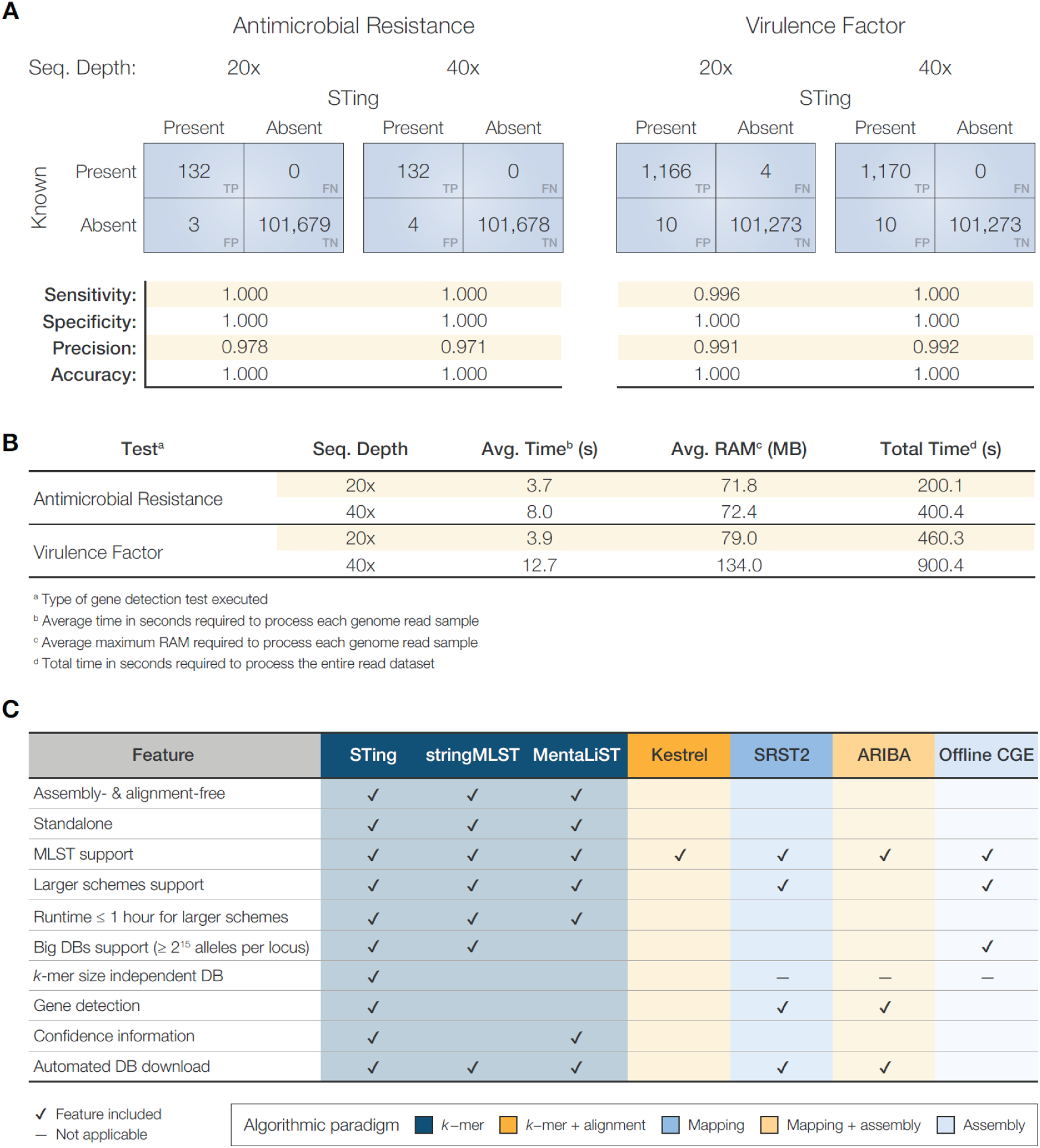
Performance comparison of STing’s Gene Detection program. STing’s Gene Detection program was run on 71 WHO designated high-priority bacterial genomes (simulated at a read depth of 20x and 40x) that contained gene annotations for 1,434 antimicrobial resistance (AMRs) and 1,443 virulence factors (VFs). (A) Confusion matrices for the detection of AMR genes from the CARD dataset, and VF genes from the VFDB dataset are shown. (B) The table demonstrates the accuracy and average runtime performance comparison of STing’s Gene Detection at each sequencing read depth. (C) Feature comparison between STing and the six applications tested for sequence typing.

STing was developed to provide turn-key solutions for NGS analysis in support of public health. Despite its lightweight computational footprint, STing is able to perform accurate and ultrafast molecular typing and gene detection. We summarize the features and utility of STing compared to related programs for genome-enabled typing in Figure 3C. In addition to its superior accuracy and performance, STing is distinguished by its streamlined algorithmic design, its broad applicability across typing schemes, its ability to support large databases, and its broad use as an automated gene detection utility.

## Data availability

Whole genome sequencing samples used for sequence typing, assemblies used for the limit of detection and multicore performance test, and genomes used for gene detection, are listed with accession numbers in the Supplementary Data.

## Code availability

The source code of STing is available at https://github.com/jordanlab/STing. The modified script implementing the Offline CGE MLST method is available at https://github.com/hspitia/binf_scripts/blob/master/run_MLST.single_thread.py.

## Online Methods

### Algorithm overview

Given an input sequence read file from a microbial isolate, STing can accurately identify the specific sequence type (ST), e.g. multilocus sequence type (MLST) or its variants, for the isolate, and what genes of interest are present in its genome. STing accomplishes these tasks by using an exact *k*-mer matching and frequency counting paradigm. STing is implemented in C++ and utilizes two libraries: the SeqAn library^10^ for the Enhanced Suffix Array (ESA)^6^ data structure and the gzstream (https://www.cs.unc.edu/Research/compgeom/gzstream/) library for working with gz files. Additionally, STing is prepackaged with an R script for visualization of the results and a Python script for downloading database sequences from PubMLST. The ESA data structure is used for *k*-mer look-up and comparison purposes. ESAs are a lexicographically sorted array-based data structure, which represent space efficient implementation of the Suffix Trees data structure. For a given set of sequences with a total length of *n* base pairs (summation of the length of all sequences), an ESA index can be constructed in linear time *O(n)*. ESAs can also be queried for *k*-mer matches (or substring matches) in linear time. Given a *k*-mer of length *k*, we can determine its presence/absence in the database in *O(k)* time and find all of its *z* occurrences in *O(k+z)* time. While Suffix Trees achieve the same time complexity for index construction and *k*-mer lookup, they take five times more storage space than ESAs. An efficient implementation of a Suffix Tree can use up to 20 bytes per input database character, whereas an equivalent ESA consumes 4 bytes per input database character. Using ESAs for *k*-mer lookup and comparison allows STing to efficiently scale with large sequence databases. The STing algorithm is divided into three steps: (1) database indexing, (2) sequence typing, and (3) gene detection (Supplementary Figure 1). Each step is described in the following sections.

### Database indexing

In this step, STing constructs an ESA index that is used during the sequence typing and gene detection modes. For sequence typing, the indexer requires a multi-fasta file with all the observed alleles in a typing scheme and an additional allelic profile file that contains combinations of allele numbers (also referred to as allelic profiles) uniquely mapped to distinct STs. The indexer constructs two ESA indices, one for the allelic sequences (allele index) and one for the profile definitions (profile index). For gene detection, the indexer requires a multi-fasta file with the gene sequences that are to be screened in the input samples. Then, the indexer constructs a single ESA index of all the gene sequences provided (gene index).

### Sequence typing

In this mode, the typer identifies the ST of a given isolate by using a gene-by-gene approach. The typer utility operates in fast or sensitive execution modes. The sequence typing step comprises six algorithmic steps: (1) read filtering, (2) *k*-mer counting, (3) candidate sequence selection, (4) depth and coverage calculation, (5) allele calling and ST prediction, and (6) reporting. In the read filtering step (1), the middle *k*-mer of each sequence is searched within the allele index database. If the middle *k*-mer is not found in the allele index, the read is discarded, otherwise the read is passed on to the next step. The size of the *k*-mer is chosen in such a way as to minimalize the possibility that using the middle *k*-mer only results in the loss of useful sequence reads (default *k*=30); users can change the *k*-mer size. In the *k*-mer counting step (2), the typer *k*-merizes each read that passed the filter matching step, and then searches each *k*-mer from the read against the allele sequence index. For each *k*-mer match in the allele index, the typer increments a *k*-mer counter for the matched alleles/loci. Once all of the reads are processed, the typer normalizes the *k*-mer frequencies by the length of the corresponding allele. In the candidate sequence selection step (3), the algorithm selects the top *N* alleles that have the maximum normalized *k*-mer frequency for each locus. For the fast execution mode, the default value of *N* is 1, and for the sensitive execution mode the default value is 3 and can be configured by the user. In the depth and coverage calculation step (4; only applicable in sensitive mode), the typer reduces the false positives by identifying regions of the candidate alleles that are not covered by any *k*-mer, and identifying any sharp valleys in the *k*-mer depth distribution across the candidate allele. This step calculates the number of *k*-mers that had a match at each base of the top *N* alleles in each locus. To speed-up this calculation, the typer constructs a smaller index consisting of only the top *N* candidate alleles, and parses the subset of reads that passed the initial *k*-mer filter (useful reads). The typer *k*-merizes the useful reads and records the location (base) of each *k*-mer in the matched allele of the smaller index. The algorithm calculates the *k*-mer depth at each base along each allele using the match start positions. The typer then looks for discontinuities in the *k*-mer depth by checking the *k*-mer depth ratio of each adjacent position. The application detects a discontinuity if the ratio is outside the range of 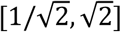 and sets the *k*-mer depth as zero for those positions. Finally, the tool calculates the allele coverage as the percentage of allele (i.e., the allele sequence length) that has a non-zero *k*-mer depth. In the allele calling and ST prediction step (5), STing generates the allelic profile and predicts the corresponding ST of the sample. For the fast mode, the allelic profile is generated from the candidate sequences selected in the previous step (step 3). For the sensitive mode, the allele with the maximum allele coverage for each locus is predicted to be the allele present within the isolate. Here, there are three special cases: (a) in the event that the allele coverage is less than 100%, the detector appends a * character to denote a possible novel allele; (b) in the event of having ties in coverage between alleles, STing calls the allele that has the most uniform *k*-mer coverage by selecting the one with the minimum *k*-mer depth standard deviation; (c) if a locus has no matching *k*-mers, the locus is assumed to be absent and an *NA* allele is assigned as its call. At this step, all the allele calls have been made and an allelic profile has been generated. A look-up operation is performed in the profile index to identify the ST corresponding to the predicted allelic profile. Finally, in the reporting step (6), STing reports the allelic profile, associated ST, and the total number of *k*-mer matches and reads processed, along with optional information about each allele: normalized counts of *k*-mer matches, coverage, and average and per-base *k*-mer depth.

### Gene detection

The algorithm for this mode is a variant of the sequence typing mode and follows the steps described above closely. The gene detection mode differs from the sequence typing mode in how it selects the candidate sequences. This mode can be divide into five conceptual steps: (1) read filtering, (2) *k*-mer counting, (3) candidate sequence selection, (4) depth and coverage calculation, and (5) reporting. In the *k*-mer filtering step (1), the detector searches the middle *k*-mer of each read within the gene index. If the *k*-mer fails to match any sequence within the index, the read is discarded, otherwise it is passed on to the next step. In the *k*-mer counting step (2), the utility proceeds to *k*-merize the read in its entirety and searches each *k*-mer in the gene index. A gene-specific *k*-mer match counter is incremented for each *k*-mer that matches the corresponding gene(s). In addition, the detector also records the start position of the *k*-mer in the matching gene(s). In the candidate sequence selection step (3), STing selects the gene sequences that have at least one *k*-mer match as probable genes present in the sample analyzed. In the depth and coverage calculation step (4), similar to the sequence typing mode, STing looks for discontinuities in the *k*-mer depth by inspecting the (a) the number of bases not covered by any k-mer, and (b) any sharp valleys within the *k*-mer distribution. Finally, in the reporting step (5), STing determines the presence/absence of genes with *k*-mer hits along with the percent sequence coverage of each gene identified in the sample. A gene is predicted to be present if its coverage is equal to or greater than a user specified threshold (default = 75%). Otherwise, the gene is predicted to be absent in the sample. STing reports the presence (reported as 1) or absence (reported as 0) of each gene with *k*-mer matches and the total number of *k*-mer matches and reads processed, along with optional information about each gene: normalized counts of *k*-mer matches, coverage, and average and per-base *k*-mer depth.

### Genomic data for sequence typing

We used 1,050 Illumina sequencing read sets of isolates from four bacterial species (*Campylobacter jejuni, Chlamydia trachomatis, Neisseria meningitidis*, and *Streptococcus pneumoniae*) retrieved from the PubMLST (https://pubmlst.org/)/EBI ENA (https://www.ebi.ac.uk/ena) database to execute the experiments (Supplementary Data). Using the isolate metadata available on PubMLST, we selected 40 samples from the four species (10 samples each) for the MLST comparative test, and 20 samples of *N. meningitidis* isolates for the larger typing schemes (rMLST and cgMLST) comparative test. We selected these two datasets trying to capture the diversity of the most common STs of each species in the PubMLST database and preferring recently sequenced isolates. For the large-scale accuracy test, we used a dataset of 1,000 *N. meningitidis* isolates.

### Computational environment

We used a machine provided with RedHat Linux SO, 24 cores, and 64 GB of RAM to perform the experiments described in this study.

### MLST comparative test design

To measure the performance of our application on the traditional seven loci MLST analysis, we compared STing (v0.24.2) in two execution modes, fast and sensitive, along with six applications able to perform sequence typing (stringMLST^4^, MentaLiST^11^, Kestrel^12^, SRST2^13^, ARIBA^14^, and Offline CGE/DTU; Supplementary Table 3). These applications can be classified into five groups depending on the strategy (algorithmic paradigm) used to predict the sequence types of whole genome sequencing data samples from bacterial isolates: *k*-mer, *k*-mer plus alignment, mapping, mapping plus local assembly, and assembly (Supplementary Table 3). For the Offline CGE/DTU application, we used the script `runMLST.py`^15^ (https://github.com/widdowquinn/scripts/blob/master/bioinformatics/run_MLST.py), an offline implementation of the original alignment-based MLST method from the Center of Genomic Epidemiology^16^. This implementation uses multithread BLAST searching for the MLST analysis, as opposed to STing, which is a single thread application. To fairly compare STing with the Offline CGE/DTU implementation, we modified the script `runMLST.py` to use only one thread for BLAST searches. For each application, we measured the accuracy in terms of the percentage of alleles correctly predicted from the total samples analyzed and the performance in terms of average run time and average peak of RAM required to analyze each of the 40 samples in the dataset. We reported the average run time and average max RAM as the average of three executions of each application per sample analyzed. Kestrel requires the generation of a *k*-mer counts file before it can be run to predict STs. For this purpose, we used the application KAnalyze^17^ (v2.0.0) with the parameters as described ^12^. We reported the average run time of Kestrel as the sum of the average times of KAnalyze and Kestrel for processing each sample and the average RAM consumption as the maximum average peak of RAM consumed by the two applications on each sample. Since the Offline CGE/DTU application requires complete assemblies to predict STs, we assembled each isolate read sample using the application SPAdes^18^ (v3.13.0) with default parameters. We reported the average runtime as the sum of the average times of SPAdes and Offline CGE/DTU to process each sample, and the average RAM consumption as the maximum average peak of RAM consumed between the two applications during the analysis of each sample. The commands used with each application tested are listed in the supplementary material (Supplementary Table 4).

### Large-scale MLST accuracy test design

To measure the accuracy of our application using the MLST scheme on a large-scale dataset, we ran STing in fast mode on 1,000 samples of *N. meningitidis*. We measured the accuracy in terms of the percentage of STs correctly predicted from the total samples analyzed, and the performance in terms of average run time and average peak RAM required to analyze each of the 1,000 samples of the dataset. We reported the average run time and average maximum RAM as the average of five executions of the application per sample analyzed.

### Limit of detection and performance on single and multicore environments test design

We evaluated the minimum sequencing depth required for correctly predicting STs on whole genome sequencing samples from bacterial isolates. We retrieved 1,306 assemblies of *Campylobacter jejuni* (n=581) and *Neisseria meningitidis* (n=725) with known MLST information from the GenBank database (https://www.ncbi.nlm.nih.gov/genbank/) (Supplementary File 1). Then, we simulated Illumina paired-end reads – HiSeq 2500, 2×150 bp, 500bp of average fragment length, with 10 as the fragment size standard deviation – from each genome at seven sequencing depths (1, 3, 5, 10, 15, 20, and 40x) using the software ART^19^ (v2.5.8). We executed STing (fast mode) over each generated sample to measure the accuracy in terms of the percentage of correct STs and alleles predicted from the total samples at each sequencing depth. We also evaluated the performance of STing in multicore environments. We executed 20 parallel instances of STing to analyze the 1,306 samples and measured the average time required to process the complete dataset at each sequencing depth.

### Large-scale sequence type schemes comparison test design

To evaluate the scalability, accuracy, and performance of our application on large-scale sequence typing schemes, we compared STing (fast and sensitive modes) on 20 samples of *N. meningitidis* against other sequence typing applications using the rMLST (loci=53) and the cgMLST (loci=1,605) schemes. We used three applications (stringMLST, SRST2, and Offline CGE) for rMLST, and three applications (stringMLST, MentaLiST, and Offline CGE) for cgMLST, which were able to execute the sequence typing analysis successfully using these larger schemes. For each application and typing scheme, we measured the accuracy in terms of the percentage of correct allele predictions from the total alleles of the tested samples and the performance in terms of the average of run time and maximum peak of RAM required to process each sample from the dataset.

### Gene detection test design

We evaluated the ability of STing to predict the presence/absence of sequences of interest in NGS read samples by detecting antimicrobial resistance (AMR) genes and virulence factor (VF) genes in simulated Illumina read datasets. We retrieved 71 assemblies from the GenBank database that correspond to 25 species listed in the World Health Organization priority list of antibiotic-resistant bacteria and tuberculosis^9^ (Supplementary Data). Then, we simulated Illumina paired-end reads – HiSeq 2500, 2×150bp, 500bp of average fragment size, with 10 as the fragment size standard deviation – from each genome at 20x and 40x sequencing depth, using the software ART. For the AMR gene detection test, we used 1,434 AMR genes available in the Comprehensive Antibiotic Resistance Database (CARD, v2.0.2)^7^. For the VF gene detection test, we used 1,443 genes from the virulence factor database (VFDB, release date 03-22-2019)^8^. In both tests, we first defined the presence/absence of each gene in each genome using BLASTn (v2.2.28+)^20^, as a ground-truth for assessing STing’s performance. To perform a fair comparison with STing’s gene detection, which is based is based on exact pattern matching, we defined a cutoff of 100% for identity and query (gene) coverage in BLASTn to consider a gene as present in a genome, i.e., if the gene is perfectly contained in the genome. Then, we built databases on STing for each gene set of interest (CADR and VFDB), and executed the respective gene detection analysis on each genome-derived read set at each sequencing depth, using a threshold of 100% for gene coverage to consider a gene as present in a sample. Finally, we evaluated the performance of detection in terms of sensitivity, specificity, precision, and accuracy, which are defined as follows:

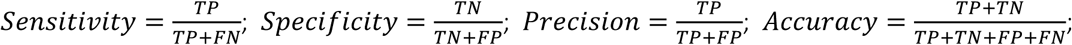

where, *TP =* true positives, *TN =* true negatives, *FP =* false positives, and *FN =* false negatives.

