## Supplementary Tables and Figures for "STing: accurate and ultrafast genomic profiling with exact sequence matches"

Supplementary Table 1. **Results of the MLST benchmarking test on a large-scale dataset.**

| Species | #Samples[b] | Depth[d] | Unique STs[c] | #Alleles[a] | Mode[e] | Correct Alleles[f] (%) | Time[g] (s) | RAM[h] (MB) | Proc. Rate[i] (MB/s) | Proc. Rate[j] (Reads/s) |
|---|---|---|---|---|---|---|---|---|---|---|
| *N. meningitidis* | 1,000 | 175.1 | 387 | 7,197 | Fast | 100.0 | 10.6 | 82.9 | 86.2 | 372,739 |
| | | | | | Sensitive | 100.0 | 11.4 | 82.9 | 80.7 | 348,760 |

[a] Total number of alleles tested
[b] Total number of samples tested
[c] Total number of unique STs in the dataset
[d] Average sequencing read depth of the dataset
[e] Execution mode of the typing tool
[f] Percentage of correctly predicted alleles
[g] Average run time in seconds per sample
[h] Average of maximum RAM usage per sample
[i] Data processing rate in megabytes per second
[j] Data processing rate in reads per second

Supplementary Table 2. **List of samples correctly predicted by STing and misannotated in PubMLST.**

| Accession[a] | Predicted ST | ST on PubMLST |
|---|---|---|
| ERR036115 | 11 | 672 |
| ERR133727 | 106 | 1087 |
| ERR133738 | 116 | 106 |
| ERR133744 | 10307 | 989 |
| ERR137178 | 144 | 102 |
| ERR310540 | 11 | 5757 |
| ERR957622 | 154 | 6697 |

[a] Accession number of the isolate in the EBI ENA database

Supplementary Table 3. **Sequence typing applications tested.**

| Application | Algorithm Type[a] | Input Type | Version | Reference[b] |
|---|---|---|---|---|
| STing | *k*-mer | Reads | 0.24.2 | This paper |
| stringMLST | *k*-mer | Reads | 0.6.1 | PMID 27605103 |
| MentaLiST | *k*-mer | Reads | 1.0.0 | PMID 29319471 |
| Kestrel | *k*-mer + alignment | Reads | 1.0.2dev1 | PMID 29186321 |
| SRST2 | Mapping | Reads | 0.2.0 | PMID 25422674 |
| ARIBA | Mapping + assembly | Reads | 2.13.3 | PMID 29177089 |
| Offline CGE | Assembly | Assembly | - | https://github.com/widdowquinn/scripts/blob/master/bioinformatics/run_MLST.py; Implementing method from PMID 22238442 |

[a] Algorithmic paradigm implemented by the tool
[b] Reference of the software application

Supplementary Table 4. **Commands used with each sequence typing software.**

| Application | Task | Command |
|---|---|---|
| STing | DB creation | indexer -c <config_file> -p <db_prefix> |
| | Sequence typing (fast) | typer -x <db_prefix> -1 <fastq_1> -2 <fastq_2> -k 30 -c -s <sample_name> |
| | Sequence typing (sensitive) | typer -x <db_prefix> -1 <fastq_1> -2 <fastq_2> --sensitive -s <sample_name> -k 30 -n 3 -c -a -d -t <out_depth_file> |
| stringMLST | DB creation | stringMLST.py --getMLST --species <species_name> -k 35 -P <db_prefix> |
| | Sequence typing | stringMLST.py --predict -1 <fastq_1> -2 <fastq_2> -k 35 -P <db_prefix> |
| MentaLiST | DB creation | mentalist build_db -k 30 --db <out_db_file> -p <profile.txt> -f <fasta_files> |
| | Sequence typing | mentalist call --db <db_file> -o <out_file> -1 <fastq_1> -2 <fastq_2> |
| Kestrel | *k*-mer counts file creation (KAnalyze) | java -Xmx3G -jar kanalyze.jar count -k 31 --countfilter=kmercount:5 --quality=10 -m ikc --minsize 15 -o <kanalyze_out_File> <fastq_1> <fastq_2> |
| | Sequence typing | java -Xmx2G -Dlogback.configurationFile=logback.xml -jar kestrelmlst.jar <allele_seqs_file> <kanalyze_out_file> > <kestrel_mlst_out_file> |
| SRST2 | DB preparation (Samtools and Bowtie2) | samtools faidx <allele_seqs_file>;<br>bowtie2-build <allele_seqs_file> <allele_seqs_file> |
| | Sequence typing | srst2 --output <out> --input_pe <fastq_1> <fastq_2> --mlst_db <allele_seqs_file> --mlst_definitions <profile_file> --mlst_delimiter '_' |
| ARIBA | DB creation | ariba pubmlstget <species_name> <out_dir> |
| | Sequence typing | ariba run <db_ref_dir> <fastq_1> <fastq_2> <out_dir> |
| Offline CGE | Assembly (SPAdes) | spades.py -1 <fastq_1> -2 <fastq_2> -o <out_dir> -t <threads> |
| | Sequence typing | run_MLST.single_thread.py -o <out_dir> -i <scheme_dir> -g <genomes_dir> -p <profile> -l <log_file> -v --force |

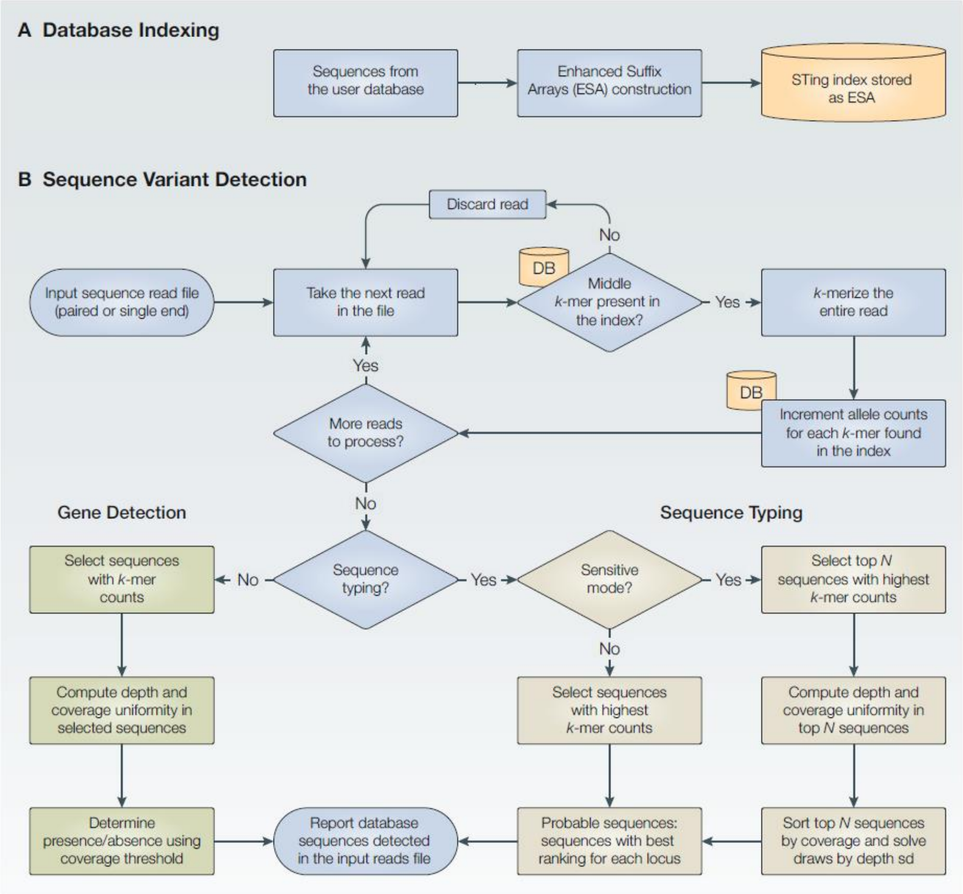


Supplementary Figure 1. **Detailed flowchart of the STing algorithm.** The STing algorithm comprises two main phases: (A) Database indexing – user supplied reference sequences (allele or gene sequences) are transformed into an Enhanced Suffix Array (ESA) index database for rapid *k*-mer search during the sequence variant detection phase; and (B) Sequence variant detection – The middle *k*-mer of each read is searched within the ESA index. If the middle *k*-mer is not found in the index, the read is discarded; otherwise, the read is passed to the next step. This filtering step allows for massively increased processing speed, since the vast majority of sequence reads in a whole genome sample do not correspond to the loci/genes found in the reference database. Reads that passed the match filter are fully *k*-merized, and each *k*-mer is searched in the ESA index. For each match in the index, a table of frequencies (*k*-mer frequency table) is updated for the matched alleles. For the sequence typing task (tan colored boxes), the top *N* alleles that have the maximum *k*-mer frequencies of each locus of the typing scheme are selected as candidate sequences to be present in the sample analyzed. If the fast mode is enabled, the value of *N* is 1, and the alleles selected for generating the allelic profile based solely in the *k*-mer frequencies. If the sensitive mode is enabled, the value of *N* is 3 by default (can be configured by the user). Then, for the sensitive mode, the allele coverage and *k*-mer depth is calculated for each of the *N* alleles in each locus. This allows for removing false positive alleles on each locus by identifying pronounced valleys in the *k*-mer distribution. Alleles are called by taking the alleles with maximum length coverage. Ties in length coverage between alleles in the same locus are solved by taking the allele with the minimum *k*-mer depth standard deviation. Allelic profiles are generated with the called alleles and a look up operation of this profile is performed in the profiles provided by the user to identify the corresponding ST for the sample. Then, the allelic profile and it corresponding ST are reported. For the gene detection task (green colored boxes), the sequences that have at least one *k*-mer match are selected as candidate sequences to be present in the sample. Then, *k*-mer depth and gene coverage is calculated for each selected gene. Genes with a length coverage greater than a threshold provided by the user (75% by default), are considered to be present in the sample analyzed. Finally, presence/absence is reported for each of the selected genes.

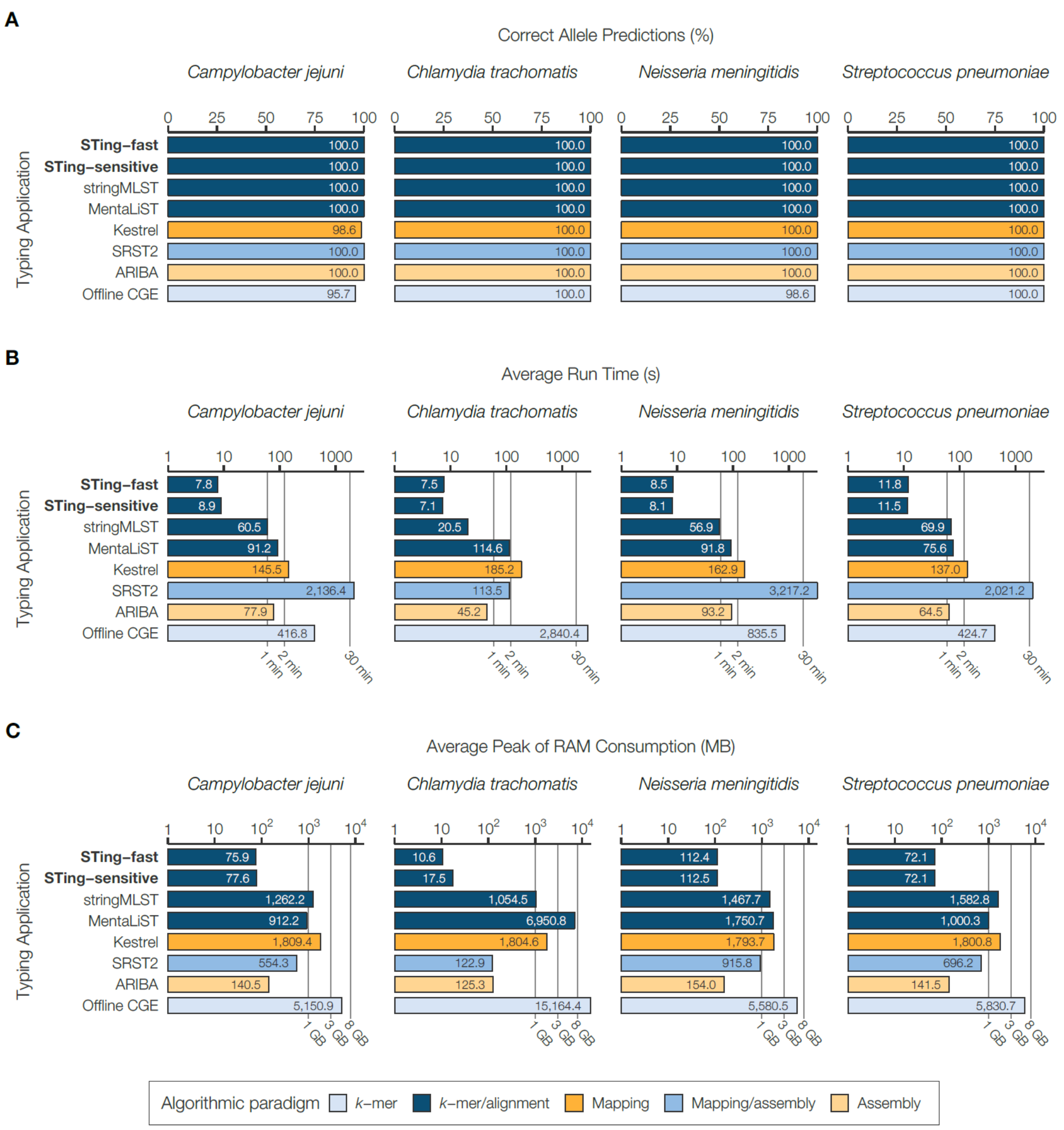


Supplementary Figure 2. **Performance comparison of STing for MLST detailed by species**. The fast and sensitive mode of STing is compared to six other contemporary typing applications to measure the accuracy and runtime performance using the traditional MLST (loci=7) on 40 samples from four bacterial species (10 samples per species): *C. jejuni*, *C. trachomatis*, *N. meningitidis*, and *S. pneumoniae*. Performance is measured in terms of (A) the percentage of correct alleles predicted, (B) the average runtime across each dataset measured in seconds (displayed in log-scale), and (C) the average peak RAM utilization across each dataset measured in megabytes (MB) for MLST, and gigabytes (GB) for rMLST and cgMLST (both displayed in log-scale). The typing applications are color-coded based on the algorithmic paradigms that they utilize for sequence typing.

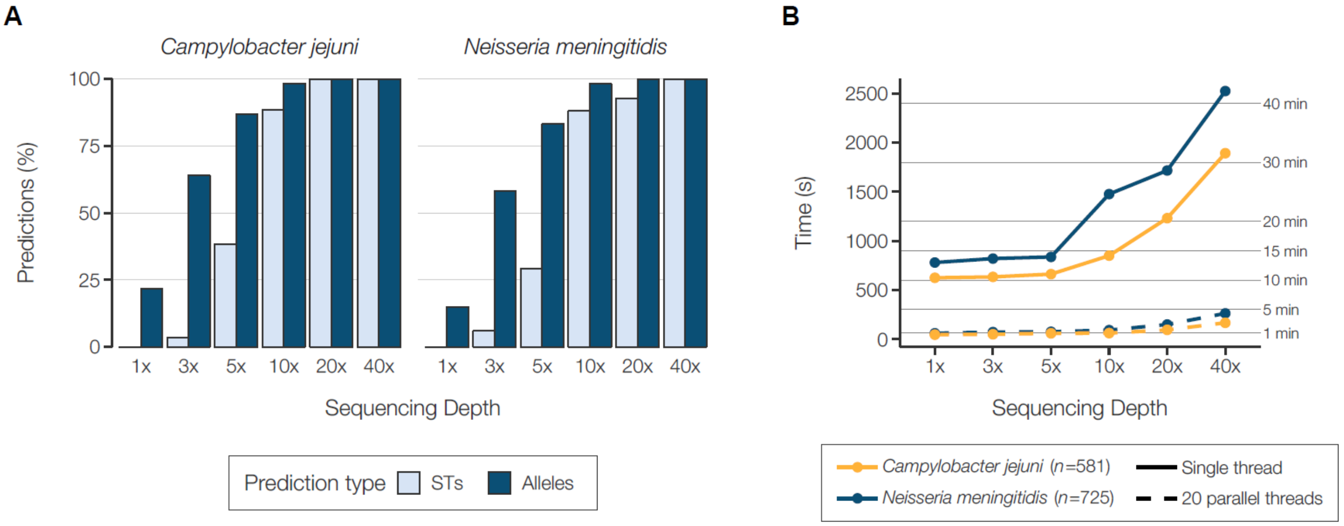


Supplementary Figure 3. **Results of the limit of detection test, and single- and multi-core performance test.** STing's typer utility was run in the fast mode over 1,306 read samples simulated at six sequencing depths (1, 3, 5, 10, 20, an 40x) from assemblies of two species *Campylobacter jejuni* (n=581) and *Neisseria meningitidis* (n=725). (A) Percentage of correct predictions in terms of STs and alleles at different sequencing depths for the two datasets. (B) Total time in seconds required to process the complete dataset for each species at different sequencing depths using a single thread or instance (solid lines) and 20 multiple threads of the typer utility.
