## Supplementary Notes for "STing: accurate and ultrafast genomic profiling with exact sequence matches"

### Pseudocode and detailed algorithms

#### 1 Database indexing

##### 1.1. Input:

Configuration file that defines relative paths to the required files:

1. Allele sequence file(s): Multi-fasta format file(s) with the sequence description being the locus name and corresponding allele number for each locus in the MLST schema.
2. Profiles file: Tab separated values file with the allelic profiles that define each ST.

##### 1.2. Output:

A set of files that defines the database:

1. Allele ESA index files: `prefix.ali`, `prefix.dat`, `prefix.ids`, `prefix.sa`, `prefix.txt.[0-N]`.
2. Profile ESA index files: `prefix.prof_idx.ali`, `prefix.prof_idx.dat`, `prefix.prof_idx.ids`, `prefix.prof_idx.sa`, `prefix.prof_idx.txt.[0-N]`,

where `prefix` is the database prefix defined by the user.

##### 1.3. General algorithm

1. Load the config file
2. Load the loci sequence files
   - 2.1. Store the allele sequences for each locus
   - 2.2. Store the allele sequence ids
3. Load the profiles file
   - 3.1. Store the profiles table
   - 3.2. Build a list of the string representation of each profile that defines an ST
4. Create and save to disk an ESA index of the list of profile string representations
5. Create and save to disk an ESA index of the allele sequences
6. Save auxiliary information
   - 6.1. Save a loci table file
   - 6.2. Save an allele-loci id pairs file
   - 6.3. Save a profiles file
   - 6.4. Save an allele sequences ids file

##### 1.4. Input files description

The database indexing stage requires a configuration file that define the location of the files that make up a typing scheme: the allele sequence files and a profile definition file. Files of the typing scheme can be downloaded from PubMLST or can be created by the user.

##### 1.4.1. Configuration file

A plain text file that contains two sections to specify the paths to the files of the typing scheme. An example of a configuration file is the following:

```
[loci]
# this is a comment line
abcZ    Neisseria_sp/abcZ.fa
adk     Neisseria_sp/adk.fa
aroE    Neisseria_sp/aroE.fa
fumC    Neisseria_sp/fumC.fa
gdh     Neisseria_sp/gdh.fa
pdhC    Neisseria_sp/pdhC.fa
pgm     Neisseria_sp/pgm.fa

[profile]
profile  Neisseria_sp/neisseria.txt
```

Each header (words between characters `[]`) define a section that contains paths to the input files. Each row of the section `[loci]` defines the allele sequence file for each locus of a typing scheme (name and file path). The section `[profile]` only has a row that specifies the profile definition file (name and file path). The configuration file must fulfill the following requirements:

- Sections headers are mandatory (`[loci]` and `[profile]`).
- Values defined in each row must be separated by TAB character.
- Blank lines and comments (lines starting with `#`) are ignored.
- File paths are relative to the location of the config file. In the example shown above, the folder `Neisseria_sp` is located at the same directory level of the configuration file.

##### 1.4.2. Allele sequence file

Each allele sequence file (one for each locus in the typing scheme) is a standard multi-fasta file with the description for each allele sequence being the locus name with the allele number. An example of the abcZ allele sequence is the following:

```
>abcZ_1
TTTGATACTGTTGCCGTAC...
>abcZ_2
TTTGATACCGTTGCCGAAA...
>abcZ_3
TTTGATACCGTTGCGAACC...
>abcZ_4
TTTGATACCGTTGCCACGT...
```

##### 1.4.3. Profile definition file

The profile definition file is a tab separated file that contains the ST and the allele profile corresponding to the ST. An example of the profile definition file is shown below:

```
ST  abcZ  adk  aroE  fumC  gdh  pdhC  pgm
1   1     3    1     1     1    1     3
2   1     3    4     7     1    1     3
3   1     3    1     1     1    23    13
4   1     3    3     1     4    2     3
```

### 2 Sequence typing

#### 2.1. Input:

1. Raw FASTQ sequencing reads (single end or paired end)
2. Database index files

#### 2.2. Output:

1. Allelic profile and associated sequence type
2. Total number of *k*-mer matches and reads processed
3. Optional information:
   a. Normalized counts of *k*-mer matches
   b. Coverage of each allele
   c. Mean *k*-mer depth
   d. Per-base *k*-mer depth file

#### 2.3. Detailed algorithm

**Algorithm 1:** STing Sequence Typing

**Input :**

Loci $L = \{l_1, l_2, \ldots, l_m\}$, $m$ is the total number of loci.

Reads $R = \{r_1, r_2, \ldots, r_n\}$, $n$ is the total number of reads.

$k$-mer size $k \leq \min(\text{length}(R))$.

Allele index $\mathcal{A}$, the generalized ESA index of all the alleles $A$ in the typing scheme, where $A = \{a_{(i,j)} \mid 1 \leq i \leq m, 1 \leq j \leq e_i\}$, $e_i$ is the total number of alleles for the locus $l_i$.

Profile index $\mathcal{P}$, the generalized ESA index of all the allelic profiles $P$ in the typing scheme, where $P = \{p_1, p_2, \ldots, p_q\}$, $q$ is the total number of observed allelic profiles.

Profile table $T = \{t_i \mid 1 \leq i \leq q\}$, where $t_i = (s_i, p_i)$, $s_i$ is the ST associated to the profile $p_i$.

**Output:**

Predicted ST and allelic profile $t' = (s_i, p_i)'$ from the read set.

```
procedure SequenceTyping(L, R, k, 𝒜, 𝒫, T)
    for each r ∈ R do                                              ▷ Read processing
        mid_kmer ← GetMidKmer(k, r)                                ▷ (i) Filtering
        if mid_kmer ∉ 𝒜 then
            continue
        freqs, hits ← []                                           ▷ (ii) k-mer counting
        K ← GetAllKmers(k, r)
        for each kmer ∈ K do
            (locus, matched_allele, hit_pos) ← Find(kmer, 𝒜)
            if matched_allele ≠ ∅ then
                freqs[locus][matched_allele]     freqs[locus][matched_allele] + 1
                hits[locus][matched_allele].add(hit_pos)
    norm_freqs ← NormalizeFreqs(freqs, 𝒢)                          ▷ Normalize by gene length
    prob_seqs ← []                                                 ▷ (iii) Selection of probable sequences
    for each locus ∈ L do
        prob_seqs.add(arg max(norm_freqs[locus]))
    (depths, coverages) ← InitDepthsAndCoverages(L, allelic_profile)
    for each allele ∈ allelic_profile do                           ▷ (iv) Depth and coverage calculation
        depths[allele] ← CalculateDepth(hits[allele])
        coverages[allele] ← CalculateCoverage(depths[allele])
    (ST, allelic_profile) ← PredictST(prob_seqs, 𝒫)                ▷ (v) ST prediction
    print ST                                                       ▷ (vi) Reporting
    for each allele ∈ allelic_profile do
        if coverages[allele] < 100 then
            print allele + "*"
        else
            print allele
```

### 3 Gene detection

#### 3.1. Input:

1. Raw FASTQ sequencing reads (single end or paired end)
2. Database index files

#### 3.2. Output:

1. Presence/absence of each gene (from the database) in the read sample
2. Total number of *k*-mer matches and reads processed
3. Optional information for each gene:
    a. Counts of *k*-mer matches
    b. Percent of *k*-mer matches from the total of *k*-mer matches
    c. Coverage
    d. Mean *k*-mer depth
    e. Per-base *k*-mer depth file

#### 3.3. Detailed algorithm

**Algorithm 2:** STing Gene Detection

**Input :**

Genes of interest $G = \{g_1, g_2, \ldots, g_m\}$.

Reads $R = \{r_1, r_2, \ldots, r_n\}$, $n$ is the total number of reads.

$k$-mer size $k \leq \min(\text{length}(R))$.

Coverage threshold $\theta$.

Gene index $\mathcal{G}$, the generalized ESA index of $G$.

**Output:**

Presence/absence predictions $P = \{(g_i, b_j) \mid g_i \in G, b_j \in B\}$, where $B = \{\text{present, absent}\}$.

```
procedure GeneDetection(R, k, θ, 𝒢)
    for each r ∈ R do                                                ▷ Read processing
        mid_kmer ← GetMidKmer(k, r)                                   ▷ (i) Filtering
        if mid_kmer ∉ 𝒜 then
            continue

        freqs, hits ← []                                              ▷ (ii) k-mer counting
        K ← GetAllKmers(k, r)
        for each kmer ∈ K do
            (matched_gene, hit_pos) ← Find(kmer, 𝒢)
            if matched_gene ≠ ∅ then
                freqs[matched_gene] ← freqs[matched_gene] + 1
                hits[matched_gene].add(hit_pos)

    prob_seqs ← []                                   ▷ (iii) Selection of probable sequences
    for each gene ∈ G do
        if freqs[gene] > 0 then
            prob_seqs.add(gene)

    (depths, coverages) ← InitDepthsAndCoverages(G, prob_seqs)

    for each gene ∈ prob_seqs do                     ▷ (iv) Depth and coverage calculation
        depths[gene] ← CalculateDepth(hits[gene])
        coverages[gene] ← CalculateCoverage(depths[gene])

    for each allele ∈ prob_seqs do                                    ▷ (v) Reporting
        if coverages[gene] ≥ θ then
            print gene + "present"
        else
            print gene + "absent"
```
